# Cell-type specific visualization and biochemical isolation of endogenous synaptic proteins in mice

**DOI:** 10.1101/363614

**Authors:** Fei Zhu, Mark O. Collins, Johan Harmse, Jyoti S. Choudhary, Seth G. N. Grant, Noboru H. Komiyama

## Abstract

In recent years, the remarkable molecular complexity of synapses has been revealed, with over 1000 proteins identified in the synapse proteome. Although it is known that different receptors and other synaptic proteins are present in different types of neurons and synapses, the extent of synapse diversity across the brain is largely unknown, mainly owing to technical limitations. Combining mouse genetics and proteomics we have previously reported highly efficient methods for purification of synaptic protein complexes under native conditions. In that approach, tandem affinity purification (TAP) tags were fused to the carboxyl terminus of PSD95 using gene targeting in mice. Here we report an approach that restricts tagging of endogenous PSD95 to cells expressing Cre recombinase. In addition, we developed a labelling strategy enabling visualization of endogenous PSD95 tagged by fluorescent proteins in Cre-expressing cells. We demonstrate the feasibility of proteomic characterisation of synapse proteomes and visualization of synapse proteins in specific cell types. We find that composition of PSD95 complexes purified from specific cell types differs from those extracted from tissues with diverse cellular composition. Therefore, these novel conditional PSD95 tagging lines will not only serve as powerful tools for precisely dissecting synapse diversity in specific subsets of regions/neuronal cells, but also provide an opportunity to better understand brain region-specific alterations associated with various psychiatric/neurological diseases. The newly developed conditional gene tagging methods can be applied to many different synaptic proteins and will thus facilitate research on the molecular complexity of synapses.

## Introduction

Information processing in the mammalian brain depends on synapse proteins. Over the past two decades, proteomic studies have revealed remarkable synapse complexity and diversity, in which biochemically purified protein complexes from the postsynaptic density were analysed and found to contain over 1000 different protein components (Peng *et al*., 2004; Collins *et al*., 2006; Trinidad *et al*., 2006; Grant, 2007; Emes *et al*., 2008; Trinidad *et al*., 2008; Distler *et al*., 2014; Uezu *et al*., 2016; Bayes *et al*., 2017; Roy *et al*., 2018). The functional diversification of synapses is often related to their molecular diversity. It has been reported that many synaptic proteins, including different subtypes of receptors, isoforms of scaffolding proteins and adhesion molecules, are differentially expressed in various brain regions (Husi *et al*., 2000; Komiyama *et al*., 2002; Lein *et al*., 2007; Micheva *et al*., 2010; Bayes *et al*., 2011; Bayes *et al*., 2012; Frank *et al*., 2016; Frank *et al*., 2017; Roy *et al*., 2018).

Despite the recent progress, we still have insufficient knowledge with regards to precisely how individual synapses within a certain type of neurons/brain region are distributed, and whether and how they are specified based on the differential molecular components. This is partly due to experimental limitations in accurately tracking and dissecting specific anatomical regions and populations of neuronal cells. It is therefore necessary to establish an experimental system that allows precise and reliable identification and characterisation of synapse diversity *in vivo*.

In recent years, gene targeting approaches to engineer the mouse genome have proven to be a crucial tool for the detailed investigation of genes of interest *in vivo*. Combined with the *Cre/loxP* recombinase system, it is possible to achieve more sophisticated genetic manipulations such as precise spatiotemporal control of gene expression, introduction of point mutations or deletions of genomic sequence. Using gene targeting, we have previously reported the generation of a constitutive PSD95-TAP tag knock-in mouse line (Fernandez *et al*., 2009), in which PSD95 (also known as PSD-95 or DLG4), an important postsynaptic scaffolding protein that plays key roles in synaptic development, protein complex assembly and plasticity, was in-frame fused with the small peptide TAP tag for highly efficient synaptic protein purifications. In the present study, we have further developed this methodology and present the generation of two lines of conditional knock-in tagging mice: PSD95-c(mCherry/eGFP) (monomeric Cherry, enhanced green fluorescent protein) and PSD95-cTAP (conditional TAP tag). In contrast to the global labelling of PSD95, here we applied a site-specific recombination strategy that allows Cre-mediated switching of fluorescent reporters that faithfully reflect endogenous expression. As a result, these genetically modified mice express red fluorescent PSD95-mCherry prior to Cre-mediated recombination and green fluorescent PSD95-EGFP after recombination, thereby allowing visualisation and distinction of PSD95 synaptic clusters in recombined and non-recombined cells. We also describe a conditional PSD95-cTAP tagging line, whereby the PSD95-TAP tag is expressed upon Cre-mediated recombination, thus allowing efficient isolation and purification of PSD95 protein complexes from recombined neurons.

Crossing these conditional tagging mice with different driver lines that express Cre recombinase under the control of specific types of neuronal promoters, we detected robust expression of PSD95-eGFP or PSD95-TAP in corresponding regions/neuronal populations. Intriguingly, we also detected two distinctive populations of PSD95 positive synaptic clusters at the CA1 and CA3 pyramidal neurons: not only different synaptic features such as synaptic density, size and punctate morphology can be observed, but also differential molecular compositions of these PSD95 complexes were revealed by proteomic analysis.

## Materials and methods

All animal procedures were carried out in accordance with the United Kingdom Animals Act (Scientific Procedures) of 1986 and approved by the British Home Office.

### Targeting vectors construction

A previously constructed intermediate targeting vector for the generation of PSD95-TAP tag mice (Fernandez *et al*., 2009) was used for tagging the C-terminus of endogenous PSD95 proteins. For generating the conditional fluorescent gene targeting vectors, a *loxP*-flanked cassette containing red fluorescent protein mCherry coding sequence (Clontech, 632523) and an FRT site-flanked neomycin resistance gene (PGK-neo) cassette were assembled into the backbone plasmid pNeoflox (Figure 1A). To generate the Cre-mediated conditional replacement of mCherry with eGFP, coding sequence for eGFP (Clontech, GenBank accession U55762) was further subcloned into the intermediate vector pNeoflox-mCherry and immediately after the FRT-PGK-neo-FRT-*loxP* sequence (Figure 1A). Both mCherry and eGFP coding cassettes were in-frame inserted just before the STOP codon of the murine *PSD95 (Dlg4)* gene using 11 amino acids encoded by nucleotide sequence for the *loxP* site as part of a flexible linker (Figure 1A). Approximately 4 kb to 6 kb upstream and downstream regions of *PSD95* last exon (with corresponding genomic sequence retrieved from BAC clones previously used in Fernandez *et al*., 2009) were cloned into the targeting vectors as 5’ and 3’ homology arms, respectively. All final targeting vectors contain a diphtheria toxin A (DT-A) fragment that allows for negative selection in embryonic stem cells.

**Figure 1.**
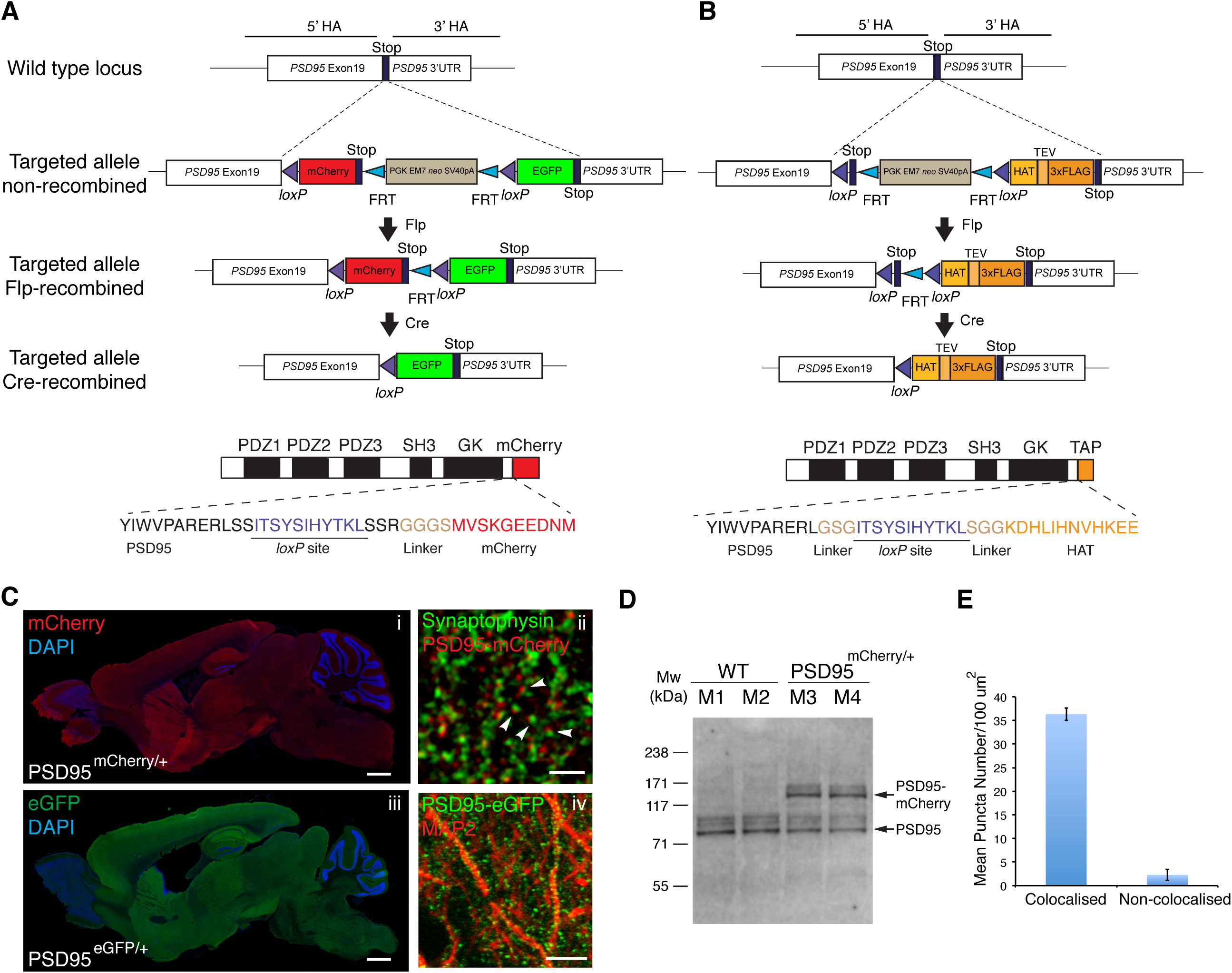
Generation of PSD95^c(mCherry/eGFP)^ and PSD95^cTAP^ knock-in mice. **(A)** Gene targeting strategy for the PSD95^c(mCherry/eGFP)^ mice. The *PSD95 (Dlg4)* allele was targeted with tandem fluorescent tag (mCherry and eGFP) coding sequence inserted at the last exon and immediately before the STOP codon. The FRT site-flanked neo gene was removed by crossing PSD95^c(mCherry/eGFP)^ mice with FLPe deleter mice. The progeny PSD95^c(mCherry/eGFP)^(without neo) mice were further bred with different Cre driver lines. Bottom panel shows domain structure of PSD95-mCherry fusion protein, which contains three PDZ, an SH3, a GK domain and the C-terminal mCherry tag (before Cre recombination). Note that a short peptide encoded by the *loxP* site and a linker sequence were inserted into the open reading frame of PSD95. **(B)** Gene targeting strategy for the PSD95^cTAP^ mice. By a similar targeting strategy, a *loxP* site-flanked STOP codon and the TAP sequence were inserted before the PSD95 STOP codon. Bottom panel shows the domain structure of PSD95-cTAP fusion protein (after Cre recombination), which includes the C-terminal-tagged TAP tag. **(C)** Ubiquitous PSD95-mCherry/eGFP expression in adult mouse brain before **(i)** and after (iii) breeding with a germline CAG-Cre driver line. Note that both PSD95-mCherry (identified by anti-mCherry antibody immunostainings, i) and PSD95-eGFP (identified by native eGFP fluorescence, iii) widely express across the brain showed similar distribution pattern. Scale bar: 0.5 mm. (ii) Fluorescence confocal image on brain sections of fluorescent knock-in PSD95^mCherry/*^ mice; PSD95-mCherry puncta (red) are located in close opposition to the anti-synaptophysin-stained pre-synaptic terminals (green)(arrowheads). Scale bar: 2 μm. (iv) Representative image of anti-MAP2 immunofluorescence staining on PSD95^eGFP/*^ brain sections. Discrete PSD95-eGFP puncta (green) were detected along the MAP2-staining neuronal processes. Scale bar: 10 μm. **(D)** Western blotting analysis of homogenate extracts from wild-type (M1, M2) and littermate heterozygous (M3, M4) PSD95^mCherry/*^ mice, using antibodies against murine PSD95. **(E)** Mean punctum number/100 μm^2^ shows that the majority of PSD95-mCherry puncta are in close opposition to (defined as ‘colocalisation’) synaptophysin-labelled pre-synaptic terminals. Error bars: mean±s.e.m; n= 3 cryosections.

The conditional PSD95-TAP targeting vectors were designed and constructed using a similar strategy to that mentioned above, in which the last exon of *PSD95* was engineered to in-frame fuse to a *loxP* site-flanked STOP codon, which is followed by a short G-S-G linker peptide coding sequence plus the TAP coding sequence, which includes a histidine-affinity tag (HAT), TEV protease cleavage site and a triple FLAG tag. Therefore, in the presence of Cre recombinase, the strategically placed STOP codon is deleted, which drives the expression of the in-frame fusion protein PSD95-TAP (Figure 1B).

Embryonic stem (ES) cell gene targeting and generation of reporter mice The targeting vectors were transfected into murine ES cells (E14 TG2a) via electroporation as previously described (Fernandez *et al*., 2009). G418 (300 μg/ml final concentration) was added to the ES cell culture 24 h after plating for positive selection. Single G418-resistant colonies were picked 5–7 days after selection.

Correctly targeted ES cell colonies were identified by long-range PCR amplification (Expand Long Template PCR system, Roche, Cat No. 11681842001) and further injected into recipient blastocysts from C57BL/6J mice. Adult male chimaeras were selected to breed with C57BL/6J wild-type female mice (The Jackson Laboratory) to produce heterozygotes, which were mated with FLP deleter mice to remove the FRT-flanked neo cassette. Genomic DNA extracted from all F1 progeny ear clips was analysed by PCR to confirm the genotype (data not shown).

Two lines of transgenic mice with region-specific expression of Cre recombinase were used to establish the conditional reporter mouse colonies: (i) Grik4-Cre expressing Cre recombinase in the CA3 pyramidal neurons of the hippocampus (Nakazawa *et al*., 2002); and (ii) Pvalb-Cre mice expressing Cre recombinase under the control of the endogenous parvalbumin *(Pvalb)* promoter (Hippenmeyer *et al*., 2005).

### Histology, fluorescence microscopy imaging

Anesthetised mice were transcardially perfused by 4% paraformaldehyde (PFA; Alfa Aesar, 30525-89-4) in 0.1 M phosphate buffer (pH 7.4). The brain was immediately post-fixed in the same fixative (4% PFA) for 3–4 h at 4°C before transferring into 30% (w/v) sucrose (in 0.1 M phosphate buffer; Sigma-Aldrich) at 4°C for at least 24 h. Coronal or sagittal sections of 18 μm thickness were cryosectioned by a cryostat (Thermo Fisher).

For some tissue sections, immunohistochemical stainings were carried out. Mounted frozen sections were thawed at room temperature and rehydrated with PBS for 10 min. After incubating sections with blocking solution 5% bovine serum albumin (BSA) in Tris-buffered saline (pH 7.5) with 0.2% Triton X-100 for 2 h at room temperature, sections were incubated with primary antibodies including anti-PSD95 (mouse monoclonal IgG1, NeuroMab, 1:250, 75-348), anti-mCherry (rabbit polyclonal, Abcam, 1:500, ab167453), anti-synaptophysin (mouse monoclonal IgG1, Millipore, 1:250, MAB5258-20UG), anti-MAP2 (rabbit polyclonal, Abcam, 1:1000, ab32454) or anti-parvalbumin (mouse monoclonal IgG1, Swant, 1:200, PV235) in a humidified chamber overnight at 4°C. After washing three times, sections were incubated with Alexa Fluor 488/546/633-conjugated secondary antibodies (Life Technologies,) for 2 h at room temperature before being mounted with mounting medium. Slides were imaged by a laser scanning confocal microscope (LSM510, Zeiss) using a 63x objective (NA 1.4) or a spinning disk confocal microscope (Andor Revolution XD system) with a 100x oil-immersion Plan-Apochromat lens (NA1.4). Z-series images were collected with an optical interval of 0.1 μm for 10–13 planes. Tile scans were stitched into larger mosaics. Images were adjusted using ImageJ (National Institutes of Health) or Photoshop software for visualisation (Adobe). For fluorescence image analysis, digital image stacks were analysed by ImageJ and in-house-designed algorithms written in ImageJ macro language. Before processing, the original images were adjusted to subtract the background fluorescence intensity. All adjustments were applied equally to all images. Images were then subjected to the algorithms for punctate object segmentation and detection. Quantitative measurements of punctum size and number were performed using an ImageJ plugin 3D object counter (Bolte & Cordelieres, 2006).

### Biochemical analysis

Whole forebrains from the constitutively TAP-tagged PSD95 line were dissected from heterozygous mice. The hippocampi were isolated from the conditional TAP-tagged PSD95 line crossed with Grik4-Cre mice. Single-step affinity purifications were carried out as described previously with minor modifications (Fernandez *et al*., 2009). In brief, brain tissue was homogenised in DOC buffer [50 mM Tris pH 9.0, 1% sodium deoxycholate, 50 mM NaF, 20 mM ZnCl, 1 mM Na3VO4, 2 mM Pefabloc SC (Roche) and 1 tablet protease inhibitor cocktail/10 ml (Roche)] and clarified as described (Grant & Husi, 2001). Extracts were incubated with Dynal beads coupled with FLAG antibody for 2 h at 4°C. The resin was washed with three cycles of 15 resin volumes of DOC buffer. Captured proteins were eluted with the same buffer containing FLAG peptide. Eluted fractions containing PSD95 were pooled, concentrated in a Vivaspin concentrator (Vivascience), reduced with DTT, alkylated with iodoacetamide and separated by one-dimensional 4–12% SDS-PAGE (NUPAGE, Invitrogen). The gel was fixed and stained with colloidal Coomassie Blue and gel lanes were cut into slices and each slice was destained and digested overnight with trypsin (Roche, trypsin modified, sequencing grade) as reported previously (Bayes *et al*., 2011).

### Mass spectrometry

Extracted peptides (three fractions per sample) were analysed by nanoLC-MS/MS on an LTQ Orbitrap Velos (Thermo Fisher) hybrid mass spectrometer equipped with a nanospray source, coupled with an Ultimate 3000 Nano/Capillary LC System (Dionex). The system was controlled by Xcalibur 2.1 (Thermo Fisher) and DCMSLink 2.08 (Dionex). Peptides were desalted on-line using a micro-precolumn cartridge (C18 Pepmap 100, LC Packings) and then separated using a 120 min RP gradient (4–32% acetonitrile/0.1% formic acid) on an EASY-Spray column, 50 cm x 75 μm ID, PepMap C18, 2 μm particles, 100 Å pore size (Thermo Fisher). The LTQ-Orbitrap Velos was operated with a cycle of one MS (in the Orbitrap) acquired at a resolution of 60,000 at m/z 400, with the ten most abundant multiple-charged (2* and higher) ions in a given chromatographic window subjected to MS/MS fragmentation in the linear ion trap. FTMS target values of 3e6 and an ion trap MSn target value of 1e4 were used and with the lock mass (445.120025) enabled. Maximum FTMS scan accumulation time of 500 ms and maximum ion trap MSn scan accumulation time of 60 ms were used. Dynamic exclusion was enabled with a repeat duration of 30 s with an exclusion list of 500 and exclusion duration of 60s.

MS data were analysed using MaxQuant (Cox & Mann, 2008) version 1.2.7.4. Data were searched against mouse UniProt sequence databases (downloaded March 2012) using the following search parameters: trypsin with a maximum of two missed cleavages, 7 ppm for MS mass tolerance, 0.5 Da for MS/MS mass tolerance, with acetyl (protein N-term) and oxidation (M) set as variable modifications and carbamidomethyl (C) as a fixed modification. A protein false discovery rate (FDR) of 0.01 and a peptide FDR of 0.01 were used for identification level cut-offs. Label-free quantification was performed using MaxQuant LFQ intensities (Cox *et al*., 2014) and statistical analysis was performed using Perseus (Tyanova *et al*., 2016). Only proteins quantified in at least three replicate purifications were included in the statistical analysis. Protein Label-Free Quantification (LFQ) intensities were log_2_ transformed, missing values were imputed and t-testing was performed (using Perseus) with correction for multiple hypothesis testing using a permutation-based FDR of 0.05 to identify proteins quantitatively different between sets of purifications.

## Results

### Generation and establishment of conditional PSD95 knock-in tagged mice

To precisely probe the expression of endogenous PSD95 protein, we applied a gene targeting-based knock-in approach in which different tags (either a TAP tag or different fluorescent protein tags with distinct excitation/emission spectrums) were in-frame fused to murine PSD95 (Figure 1A,B). Previous findings have shown that the *PSD95* gene has multiple isoforms with alternatively spliced N-terminals but all shared a common C-terminus (Bence *et al*., 2005); therefore, these tags were inserted into the open reading frame and immediately before the STOP codon (Figure 1A,B). To maintain the 'in-frame' insertion of the reporter cassettes before and after Cre-mediated recombination, we designed two similar targeting vectors for the fluorescent or TAP tagging, in which the mCherry coding sequence or a STOP codon, together with the FRT-flanked selection marker cassette, were placed immediately after the first *loxP* site, which encodes 11 amino acids, plus a short glycine-rich linker sequence (G-G-G-S or G-S-G for the fluorescent or TAP tag, respectively) (Figure 1A,B). Following excision of the mCherry or STOP codon sequences that reside between the two *loxP* sites, the resulting targeted allele will convert to the coding sequence for eGFP or TAP peptide, which again follows an 'in-frame' *loxP* site together with the short glycine-rich linker sequence (Figure 1A and materials and methods).

Over 200 G418-resistant ES cell clones were screened, and correctly targeted clones were identified by long-range PCR using a 3' external genomic reverse primer to validate the integrity of the targeted allele. Clones were selected for blastocyst injection to generate chimaeras for either the conditional fluorescent tagging or TAP tagging mouse lines. Germline transmitted mice from chimaeras were identified by PCR genotyping using specific primer sets. F1 heterozygous offspring were bred with FLPe deleter mice, expressing FLP recombinase, to remove the neo cassette flanked by two FRT sites (Figure 1A,B). Heterozygous progeny (without the PGK neo cassette) were used for further breedings with either germline-expressing CAG-Cre or cell type-specific Cre driver lines.

To confirm that introducing fluorescent tags into the endogenous gene locus did not alter the expression levels of endogenous PSD95, we performed western blot analysis using protein extracts of homogenates from heterozygous PSD95^mCherry/*^ and littermate wild-type mice. Antibodies against murine PSD95 showed two bands from the heterozygous sample group, with the ~130 kDa and ~100 kDa bands being of similar intensity (Figure 1D).

In the absence of Cre, heterozygous PSD95^c(mCherry/eGFP)/*^ mice carry a PSD95-mCherry fusion allele, whereas the eGFP cassette was prevented from transcribing due to the downstream position of the STOP codon (Figure 1A). Therefore, brain sections from these mice should only be labelled with PSD95-mCherry. Fluorescence microscopy of fixed sagittal sections from adult PSD95^c(mCherry/eGFP)/*^ mice revealed that prominent mCherry expression can be observed among various brain regions (Figure 1C). Higher fluorescence intensity levels of PSD95-mCherry were detected in the forebrain, including cerebral cortex and hippocampus, whereas weaker fluorescence was detected in other regions including the midbrain, cerebellum and medulla oblongata. Confocal images revealed that most of the PSD95-mCherry puncta were situated in close opposition to the synaptophysin-immunoreactive pre-synaptic terminals, thus confirming the synaptic localisation of PSD95-mCherry fusion protein (Figure 1Ci, Cii and Figure 1E). This pattern strongly resembles previous immunohistochemical findings using a specific PSD95 antibody (Fukaya & Watanabe, 2000).

Upon Cre recombination, the *loxP* site-flanked region (including mCherry coding sequence, PGK neo cassette and the following STOP codon) was excised and the eGFP coding cassette brought immediately downstream of the *PSD95*, creating a PSD95-eGFP fusion allele (Figure 1A). To test whether the targeted allele can be converted from PSD95-mCherry to PSD95-eGFP, we crossed a male heterozygous PSD95^c(mCherry/eGFP)/*^ mouse with female CAG-Cre transgenic mice (Sakai & Miyazaki, 1997). CAG-Cre is capable of ubiquitous deletion of *loxP* site-flanked gene sequence in the germline cells. As a result, brain sections from the progeny of these breedings should be labelled with the eGFP fluorescent reporter tag. As expected, breeding with the CAG-Cre deleter line resulted in heterozygous mice expressing prominent PSD95-eGFP across the brain (Figure 1Ciii, Civ).

### Neuronal cell type-specific expression of PSD95 positive synapses in the brain

To further examine whether the targeted allele is responsive to a tissue/cell-type specific conversion, conditional heterozygous PSD95^c(mCherry/eGFP)/*^ mice were bred with Grik4-Cre or Pvalb-Cre transgenic mice, respectively. Grik4-Cre mice show Cre activity in nearly all pyramidal neurons of the hippocampal CA3 area (Akashi *et al*., 2009), whereas Pvalb-Cre mice express Cre recombinase in parvalbumin-positive neurons (Hippenmeyer *et al*., 2005). As described in Figure 2A, brain cryosections from double-transgenic PSD95^c(mCherry/eGFP)/*^; Grik4-Cre mice showed CA3 region-specific expression of PSD95-eGFP, in contrast to the unrecombined PSD95-mCherry expression in other hippocampal regions. Interestingly, higher magnification views revealed distinctive PSD95^*^ synapse populations among different hippocampal subregions: in contrast to the densely packed PSD95-mCherry puncta in the CA1 region, the PSD95-eGFP puncta in the CA3 area (stratum lucidum) appeared to be larger and more sparsely distributed (Figure 2A). Quantification of PSD95 punctum density showed a significant difference between CA1 (~120/100 μm^2^) and CA3 (~40/100 μm^2^) subregions (Figure 2B). The punctum size of PSD95 in these regions also differs (0.015 μm^3^ in CA1 versus 0.023 μm^3^ in CA3) (Figure 2B). Taken together, these data suggest that distinctive PSD95^*^ synaptic puncta occur in different subregions of the hippocampus.

**Figure 2.**
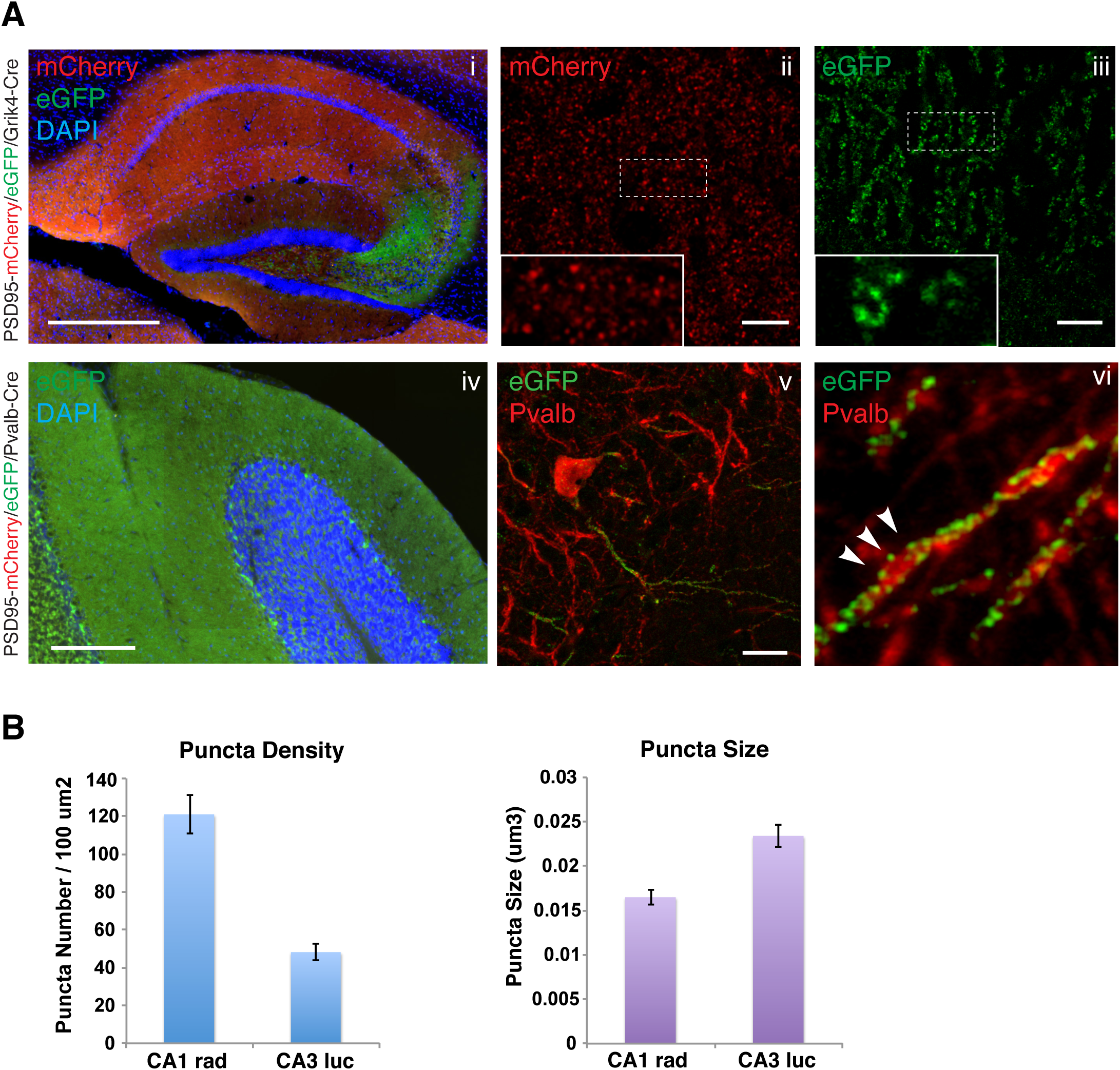
Fluorescence imaging of conditional PSD95 tagging mice reveals differential PSD95 synapses in different populations of neurons. **(A)** Fluorescence imaging in the hippocampal regions of the PSD95^c(mCherry/eGFP)^; Grik4-Cre mice. (i) Low-magnification view in the hippocampal region. Note that the CA3 subregion shows pronounced native eGFP fluorescence compared with mCherry red fluorescence in other hippocampal subregions. Scale bar: 0.5 mm. (ii,iii) Representative direct fluorescence images of CA1 and CA3 subregions demonstrating distinctive PSD95-mCherry (ii) or PSD95-eGFP (iii) punctum pattern in stratum lucidum and stratum radiatum, respectively. Insets are 5x magnifications of the boxed regions in the main images. Scale bars: 10 μm. (iv) Low-magnification micrograph of the cerebellum in PSD95^c(mCherry/eGFP)^; Pvalb-Cre mice. Native eGFP fluorescence was detected in the molecular layer of the cerebellum, as well as the pinceau termini of basket cells. Scale bar: 0.5 mm. **(v)** Representative image of CA3 Pvalb-positive neurons, labelled by anti-PV235 staining (red) together with direct GFP fluorescence of PSD95-eGFP (green). Scale bar: 10 μm. (vi) 5x magnification of region from v. Discrete PSD95-eGFP puncta were detected along the Pvalb-positive neuronal processes (arrowheads). **(B)** Quantification of PSD95 punctum density and size in CA1 and CA3 subregions of the hippocampus. Error bars: mean±s.e.m; n= 2 mice.

In addition, we also examined compound PSD95^c(mCherry/eGFP)/*^; Pvalb-Cre transgenic mice. Parvalbumin-positive cells were visualised by antibody staining. As shown in Figure 2A, discrete PSD95 puncta were observed distributed along the parvalbumin^*^ neuropil, suggesting a fraction of PSD95^*^ excitatory synapses identified in the hippocampal parvalbumin^*^ cells.

### Proteomic analysis of PSD95-cTAP in the hippocampus versus the CA3-subfield

Although we have observed distinctive features of PSD95^*^ synaptic puncta from CA1 and CA3 pyramidal cells of the hippocampus via imaging techniques, it remains unknown whether and how these PSD95 complexes differ at the molecular level.

To address this issue, we performed cell type-specific tagging of PSD95 and characterised the composition of its protein complexes using a high-affinity FLAG purification and analysis of the components using mass spectrometry.

Mice carrying the conditional PSD95-TAP tag alleles (PSD95^cTAP^) were crossed with the Grik4-Cre line in which Cre recombinase was expressed in CA3 of the hippocampus. We checked the expression of tagged PSD95 protein in the lysates from the each of the double-transgenic lines. No leaky expression of the tag could be detected in the cortex (Cx), hippocampus (Hp), or striatum (Str) of the PSD95^cTAP/*^ mice (Figure 3A). We could detect FLAG tagged PSD95 protein in the brain lysate from double-transgenic mice (PSD95^cTAP^; Grik4-Cre) (Figure 3A and 3B).

**Figure 3.**
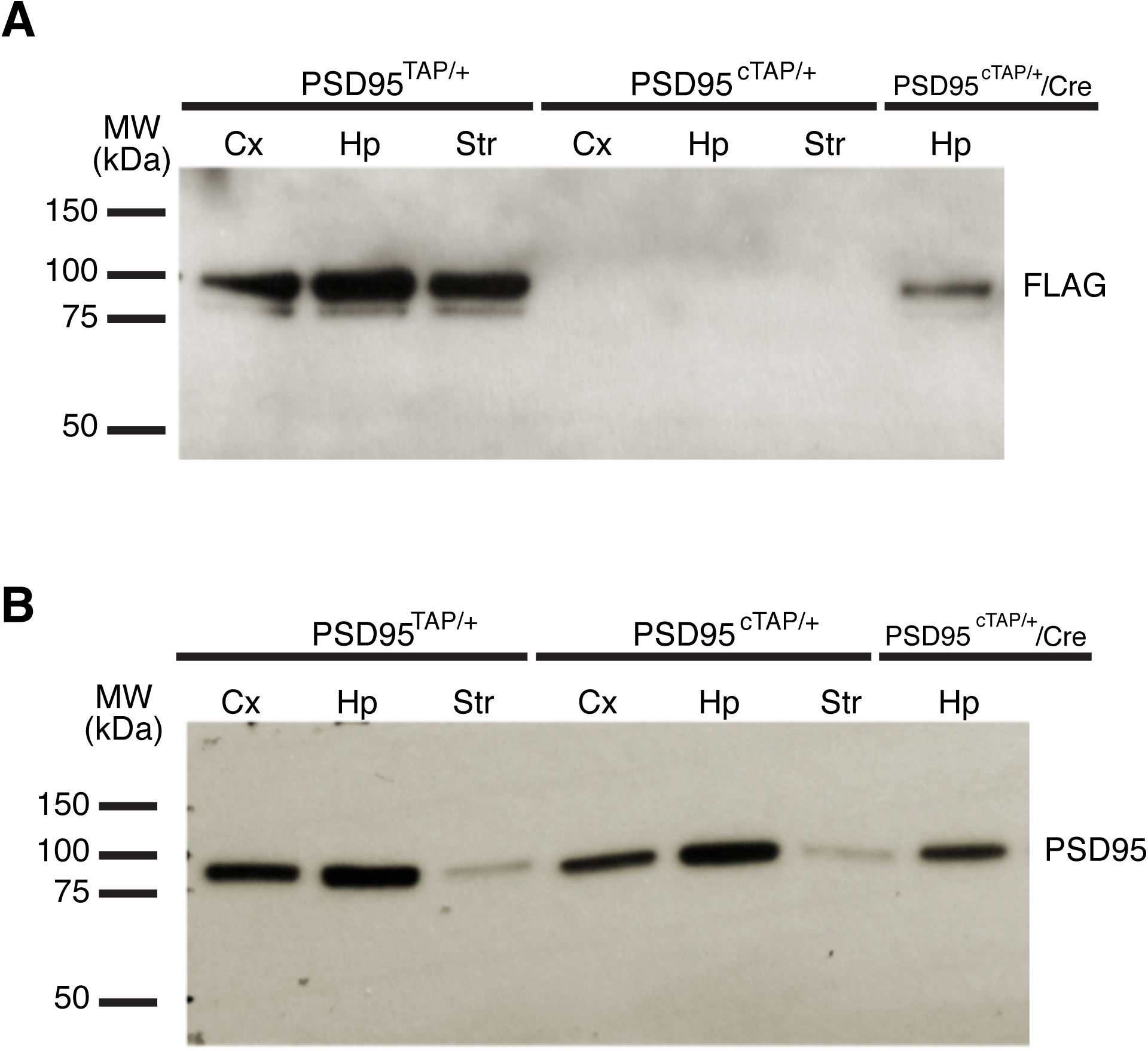
Proteomic analysis of conditional PSD95 TAP-tagged mice reveals differentially interacting proteins in specific brain subregions. **(A)** Lysates from the conditionally TAP-tagged mice were probed for FLAG to confirm expression of the tag. Expression was detected in PSD95^cTAP/*^;Grik4-Cre lines (PSD95^cTAP/*^/Cre). The lysates were taken from the hippocampi of the Grik4-Cre mice. No leaky expression of the tag could be detected in the cortex (Cx), hippocampus (Hp) or striatum (Str) of the PSD95^cTAP/*^ mice. PSD95^TAP/*^ mice were used as a positive control, and the tag was detected in all tested brain regions. **(B)** PSD95 expression was detected in all samples.

In order to generate a robust set of PSD95 interactors in the hippocampus (previous datasets were generated from forebrain tissue) (Fernandez *et al*., 2009), we performed four replicate purifications from mouse hippocampal tissue expressing the constitutively expressed TAP-tagged version of PSD95 (Fernandez *et al*., 2009) and three replicate control purifications using wild-type mice and performed protein/peptide identification and label-free quantification using MaxQuant (Cox & Mann, 2008). To define a set of proteins that were enriched in constitutively expressed PSD95-TAP purifications compared with control purifications, *t*-testing was performed (using Perseus) on LFQ protein intensities with correction for multiple hypothesis testing using a permutation-based FDR of 0.05. Of the 403 proteins quantified in at least 3 replicates, 103 proteins were significantly enriched in PSD95-TAP purifications compared to controls (Figure 4A, Table S1), representing a dataset of quantitatively enriched proteins associated with constitutively expressed TAP-tagged PSD95 in the hippocampus.

**Figure 4.**
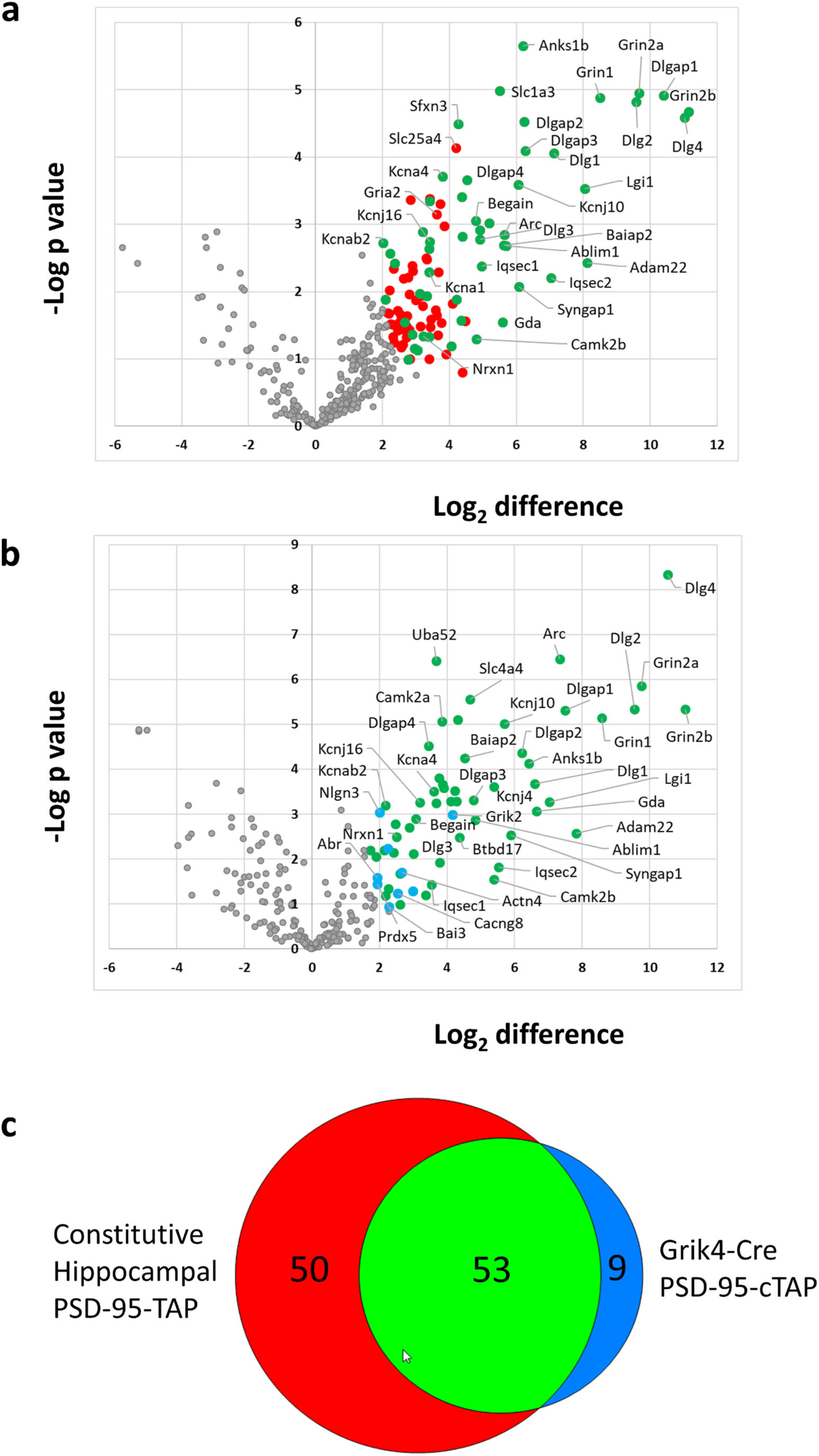
Proteomic analysis of PSD95-TAP/Grik4-Cre complexes reveals a subset of the hippocampal PSD-95 interactome. The composition of PSD-95 complexes isolated from constitutive PSD95-TAP and PSD95-cTAP/Grik4-Cre mice was compared to control purifications using label-free quantitative mass spectrometry. **(A)** Volcano plot of proteins enriched in constitutive PSD95-TAP purifications from hippocampal tissue versus wild type controls. 103 proteins were significantly enriched (Permutation-based FDR) (red and green) and 53 of these were also enriched in PSD95-cTAP/Grik4-Cre purifications (green). **(B)** Volcano plot of proteins enriched in PSD95-cTAP/Grik4-Cre purifications from hippocampal tissue versus wild type controls. 62 proteins were significantly enriched (Permutation-based FDR) (green and blue), 9 of which were not enriched in constitutive PSD95-TAP purifications from hippocampal tissue (blue). **(C)** Venn diagram of proteins enriched in constitutive hippocampal PSD95 complexes and PSD95-cTAP/Grik4-Cre complexes compared to hippocampal tissue controls. CA3 restricted (Grik4-Cre) PSD95 complexes are a subset of the total hippocampal PSD95 interactome.

Next we purified PSD95 complexes from hippocampi dissected from the PSD95^cTAP^; Grik4-Cre double-transgenic line (see materials and methods). Four replicate CA3 purifications were analysed by mass spectrometry and we defined a set of proteins that were enriched in purifications from mice with CA3 specific expression of PSD95-TAP by comparison with data from control purifications from hippocampal tissue of wild type mice. In total, of 234 proteins quantified in at least 3 replicates, 62 proteins were significantly enriched in purifications from Grik4-Cre mice representing a dataset of quantitatively enriched proteins associated with TAP-tagged PSD95 expressed in the CA3 region of the hippocampus (Figure 4B, Table S2).

Comparison of the constitutive hippocampal and CA3 specific PSD95-cTAP datasets reveal an overlap of 53 proteins with 9 that are specific to the CA3 subfield (Figure 4C) and 50 that are specific to the constitutive hippocampal dataset. The CA3 dataset contains PSD95 complex components that are mostly (90%) independently identified as complex components in the hippocampus of the constitutively expressed PSD95-TAP mouse line. Importantly, this CA3-restricted set represents just over half (51%) of the components of constitutive hippocampal dataset indicating that this approach allows the identification of brain region specific PSD95 complexes. These results demonstrate proof of concept that this tagging approach allows the purification and characterisation of synaptic protein complexes from a specific region of the hippocampus.

## Discussion

The findings reported here provide proof-of-principle for applications in conditional PSD95 labelling for both visualisation and biochemical analysis in the mouse brain. PSD95 is one of the most abundant scaffolding proteins, is widely expressed in many different types of neuronal cells in the brain and forms large macromolecular complexes with neurotransmitter receptors, various ion channels, cell adhesion and cytoskeletal proteins and signalling molecules (Husi *et al*., 2000; Husi & Grant, 2001; Collins *et al*., 2006; Fernandez *et al*., 2009; Frank *et al*., 2016; Frank *et al*., 2017). These complexes play essential roles in synaptic function, and mutations in the genes encoding the components of these complexes are implicated in human psychiatric disorders (Bayes *et al*., 2011; Kaizuka & Takumi, 2018). However, the molecular nature and heterogeneity of these complexes in different brain regions, in different neuronal types or at different developmental stages are not well understood. Therefore, it is desirable to develop an efficient approach by which the reporter transgene can be activated in a temporally and spatially regulated fashion in specific cell lineage or tissue.

Here, we present two conditional PSD95 reporter mouse lines generated by the gene targeting approach. These mouse lines allow not only simultaneous visualisation of distinctive PSD95* synaptic puncta in different populations of neurons, but also facilitate quantitative measurement and analysis of PSD95-containing synaptic protein complexes (Frank *et al*., 2016) by biochemical/proteomic methods.

A major concern with current technical efforts in molecular mapping/imaging of the synapse is the use of antibodies that may lack sufficient specificity. Another technical difficulty concerns the accessibility of antibody into the protein-dense PSD structure. In a study carried out by Fukaya and Watanabe (Fukaya & Watanabe, 2000), tissue sections were required to be processed with carefully titrated pepsin treatment before achieving specific labelling of PSD95, thus demanding substantial investment of time and efforts. Furthermore, antibody cross-reactivity, especially when detecting closely related protein paralogues, also needs to be carefully addressed.

Using fluorescent proteins genetically tagged to target protein could overcome the above limitations. In contrast to conventional immunofluorescence staining or biochemical methods mostly relying on antibodies, genetically tagging provides easy access to direct visualisation or biochemical isolation/purification of tagged protein, thus avoiding complicated processes of tissue treatment as well as antibody characterisation, a labour-intensive step to verify specificity.

It is also recognised that quantifying and comparing immunostaining signals between different brain sections is often difficult and unreliable, even using the same dilution of antibodies and staining conditions. In contrast to this general problem of quantification, we found that fluorescent signals between different slices prepared from different individual PSD95-eGFP knock-in mice were remarkably consistent and reproducible. This has allowed us to survey and numerically quantify PSD95-eGFP puncta in terms of their number, size and intensity at the whole brain scale. (Zhu *et al*., ‘Architecture of the mouse brain synaptome’, Neuron, in press, 2018). In addition, the genetic fluorescent labelling also makes it possible for live imaging of brain tissues or in the intact animal to study the molecular dynamics of PSD95 (Wegner *et al*., 2018).

Compared with other genetically labelling methods such as BAC transgenesis, which may sometimes fail to report the expression and localisation of target protein properly due to the saturation of targeting machinery or overexpression of labelled proteins, the gene targeting method enables tags to be integrated into a specific genomic locus, thus avoiding these problems. Several lines of evidence have shown that a physiological expression level of PSD95 is crucial for maintaining the normal number, size and morphology of spine synapses.

Overexpression of PSD95 modifies synaptic transmission and impairs synaptic plasticity (El-Husseini *et al*., 2000; Ehrlich & Malinow, 2004; Nikonenko *et al*., 2008). To precisely probe the expression of PSD95 and keep to a minimum the functional perturbation of the endogenous protein, we decided to apply the gene targeting-based approach. Compared with other transient transfection methods, the locus-specific in-frame insertion of the tag faithfully follows the endogenous expression of PSD95, thus avoiding the overexpression problem. Further, in contrast to other genetic labelling techniques such as insertion of the *lacZ* reporter gene, which allows quick but rather general evaluation of expression pattern at a cellular level (Porter *et al*., 2005), our PSD95 labelling enables reliable recapitulation of endogenous expression and precise protein localisation at the subcellular level.

One potential limitation of constitutively GFP tagged PSD95 knock-in is difficulties in probing the expression of tagged protein in a specific subset of cells. The conditional labellings of PSD95 in this study not only allow precise visualisation, but also faithful proteomic capture and analysis of PSD95 molecular complexes in specific subset of excitatory synapses.

Combined with appropriate Cre driver mice, these reporter lines offer an opportunity to readily label endogenous PSD95 in selected subpopulation of neuronal cells/tissues. For example, within the compound conditional fluorescent reporter line, two distinctive populations of PSD95 synaptic puncta from CA1 and CA3 pyramidal neurons identified by spectrally different fluorescent proteins were detected simultaneously in the fluorescently tagged mice (Figure 2B). Therefore, differential synaptic properties (synaptic density, size and morphology) can be readily compared and characterised at the single-synapse level. Although some difficulties remain in terms of direct comparison of fluorescence intensity between two different species of fluorescent protein, the other conditional PSD95-cTAP tagging line provides a key complementary approach for this issue.

In this study, we have also successfully isolated and purified PSD95 protein complexes from different brain regions using conditional affinity tagging methods. Our results suggest that there might be differential interactions in the PSD95 complexes in different brain regions. We have detected differentially interacting proteins by comparing datasets from the hippocampus and the CA3 subfield of the hippocampus, all these CA3 (PSD95-cTAP; Grik4 Cre) enriched proteins (Abr, Actn4, Bai3, Cacng8, Grik2, Hist1h2al, Mbp, Nlgn3 and Prdx5) have been reported to have important functions in brain that include roles for synapse formation, synaptic plasticity and psychiatric disorder (Walikonis *et al*., 2001; Oh *et al*., 2010; Lanoue *et al*., 2013; Kalinowska *et al*., 2015; Sigoillot *et al*., 2015; Duman *et al*., 2016; Ge *et al*., 2016; Martinelli *et al*., 2016; Malty *et al*., 2017; Park *et al*., 2017; Yamagata *et al*., 2017). This might implicate that different types of PSD95 complexes exist in different brain regions with distinct functions.

Functional importance of these enriched proteins in the PSD95 complexes in the specific brain area can be further elucidated by using mouse lines with conditional/floxed allele for these genes crossed with same Cre driver line (e.g. Grik4 Cre).

We previously reported the expression pattern of PSD95 and SynGAP, a major interactor of PSD95, in different brain regions and at different developmental stages (Porter *et al*., 2005). That study revealed remarkable differences in the expression pattern of these two PSD proteins at the cellular level using *lacZ* reporter knock-in mouse lines. These results indicated that the composition of PSD95 macromolecular complexes may show a high degree of heterogeneity in different neuronal cells. This view is consistent with our current finding showing that the molecular composition of PSD95 complexes in the hippocampal CA3 neurons differed from that in the rest of the forebrain. Also, our recent study combining genetics and proteomics demonstrated that there are many different types of PSD95 subcomplexes in the forebrain (Frank *et* al., 2016), and our new method could help to elucidate further details of these different complexes. Using standardised proteomic analysis methods, this conditional tagging mouse line will enable us to accurately quantify and characterise the various PSD95 supramolecular complexes in virtually any lineage of neuronal cells/tissues via different Cre driver lines.

In the future we can further genetically engineer current reporter mouse lines (i.e. via the CRISPR/Cas system) to incorporate additional fluorescent reporter gene cassette designed to follow a multicistronic element in order to visualise target neuronal morphology. As a result, both PSD95 and neuronal processes can be observed simultaneously upon Cre-mediated recombination. Therefore, this type of reporter line will be a valuable tool in making useful connections between the data generated by on-going connectome activity and molecular synaptome mapping.

Taken together, these novel conditional PSD95 reporter mouse lines will serve as a powerful tool for detailed dissection of synapse diversity in any selected subset population of neuronal cells/tissues via optical imaging or proteomic techniques. These lines can be also used in many other research fields, such as generating transgenic animals by crossing with various neurological/psychiatric disease mouse models. Our conditional tagging method can be applied to any protein complexes expressed in different tissues and cell types in the mouse to study the distinct composition and distribution of multiprotein complexes.

## Acknowledgements

We wish to thank David Fricker, Ellie Tucker and Kathryn Elsegood for technical assistances; FZ, JH, SGNG and NHK received research funding from the Wellcome Trust.

## Author contributions

NHK designed the study. SGNG and JSC provided research facilities and tools. FZ, JH and MOC performed the experiments. FZ, MOC and NHK analysed and interpreted the data. FZ, MOC and NHK wrote the manuscript.

## Data accessibility

Supporting data are available on request.

## Abbreviations

eGFP: enhanced green fluorescent protein
mCherry: monomeric Cherry fluorescent protein
TAP: tandem affinity purification
PSD: post-synaptic density

